# Flagellar stators stimulate c-di-GMP production by *Pseudomonas aeruginosa*

**DOI:** 10.1101/483339

**Authors:** Amy E. Baker, Shanice S. Webster, Andreas Diepold, Sherry L. Kuchma, Eric Bordeleau, Judith P. Armitage, George A. O’Toole

**Affiliations:** Department of Microbiology and Immunology, Geisel School of Medicine at Dartmouth; Department of Biochemistry, University of Oxford, Oxford, United Kingdom; Current Address: Department of Ecophysiology, Max Planck Institute for Terrestrial Microbiology, Marburg, Germany

**Keywords:** c-di-GMP, stators, flagella, motility, biofilm, *Pseudomonas aeruginosa*

## Abstract

Flagellar motility is critical for surface attachment and biofilm formation in many bacteria. A key regulator of flagellar motility in *Pseudomonas aeruginosa* and other microbes is cyclic diguanylate (c-di-GMP). High levels of this second messenger repress motility and stimulate biofilm formation. C-di-GMP levels regulate motility in *P. aeruginosa* in part by influencing the localization of its two flagellar stator sets, MotAB and MotCD. Here we show that just as c-di-GMP can influence the stators, stators can impact c-di-GMP levels. We demonstrate that the swarming motility-driving stator MotC physically interacts with the transmembrane region of the diguanylate cyclase SadC. Furthermore, we demonstrate that this interaction is capable of stimulating SadC activity. We propose a model by which the MotCD stator set interacts with SadC to stimulate c-di-GMP production in conditions not permissive to motility. This regulation implies a positive feedback loop in which c-di-GMP signaling events cause MotCD stators to disengage from the motor; then disengaged stators stimulate c-di-GMP production to reinforce a biofilm mode of growth. Our studies help define the bidirectional interactions between c-di-GMP and the motility machinery.

**Importance.:** The ability of bacterial cells to control motility during early steps in biofilm formation is critical for the transition to a non-motile, biofilm lifestyle. Recent studies have clearly demonstrated the ability of c-di-GMP to control motility via a number of mechanisms, including through controlling transcription of motility-related genes and modulating motor function. Here we provide evidence that motor components can in turn impact c-di-GMP levels. We propose that communication between motor components and c-di-GMP synthesis machinery allows the cell to have a robust and sensitive switching mechanism to control motility during early events in biofilm formation.

## Introduction

Two key features in biofilm formation, motility repression and exopolysaccharide (EPS) production, are controlled by the second messenger cyclic diguanylate (c-di-GMP). Intracellular levels of c-di-GMP are governed by diguanylate cyclases (DGCs) that synthesize c-di-GMP from GTP and phosphodiesterases (PDEs) that hydrolyze c-di-GMP to an inactive form (1, 2). Two DGCs in *P. aeruginosa* PA14 are critical for controlling biofilm formation: SadC and RoeA. SadC controls flagellar motility (3) and RoeA controls production of the EPS known as Pel (4). Like many bacterial species, *P. aeruginosa* has multiple DGCs and PDEs, and it is not well understood how the activity of these enzymes are coordinated to achieve signaling specificity.

Flagellar motility in *P. aeruginosa* is ultimately controlled by the flagellar motor, which generates torque from two proton-powered stator sets, MotAB and MotCD (5, 6). Only one of these stator sets, MotCD, can support swarming motility. In *P. aeruginosa*, swarming motility allows movement across a semisolid agar surface, and requires a functional flagellum and production of rhamnolipid surfactants (7). Previous studies have shown that c-di-GMP impacts stator localization, and thus flagellar motor function, by decreasing polar localization (and likely motor engagement) of MotCD (8). Furthermore, we showed that the c-di-GMP effector protein, FlgZ, physically interacts with MotC in a c-di-GMP-dependent manner. We proposed that FlgZ-MotC interactions resulted in displacement of the MotC stator from the motor, thereby reducing swarming motility (9).

Here we describe a role for the flagellar stators in controlling c-di-GMP level. We demonstrate an interaction between the DGC SadC and the stator protein MotC via SadC’s transmembrane domain. We provide evidence that this interaction stimulates SadC activity, resulting in the increased production of c-di-GMP. We propose that removal of MotC from the motor by the FlgZ:c-di-GMP effector allows enhanced opportunity for MotC and SadC to interact, resulting in a further enhancement of c-di-GMP production, and ultimately resulting in a positive feedback loop of second messenger production that we predict would effectively and rapidly reduce swarming motility.

## Results

### Loss of MotAB stators increases polar localization of GFP-MotD

We have previously reported that mutating the phosphodiesterase BifA leads to an increase in intracellular c-di-GMP and swarming motility repression in *P. aeruginosa* (10). We also demonstrated that loss of the MotAB stator restores swarming motility to the Δ*bifA* mutant, indicating that MotAB participates in c-di-GMP-mediated swarming repression (8). We hypothesized that both stator sets of *P. aeruginosa*, MotAB and MotCD, compete for engagement with the flagellar motor and that loss of MotAB allows for increased incorporation of MotCD at the motor. Since MotCD is necessary for swarming motility, increased motor engagement by MotCD would lead to increased swarming.

To test the hypothesis that localization of MotCD is altered upon loss of MotAB in the Δ*bifA* Δ*motAB* mutant, we replaced the *motD* gene at its native chromosomal locus with *gfp-motD*. We previously reported that this GFP fusion does not interfere with MotD’s function in swarming motility (8). Consistent with previous findings (8), we found that a significantly lower percentage of Δ*bifA* mutant cells have GFP-MotD polar puncta compared to wild-type (WT) cells (Figure 1A,B). In the Δ*bifA* Δ*motAB* mutant, we observed that GFP-MotD polar localization increased significantly compared to the Δ*bifA* mutant (Figure 1A,B). This finding supports our hypothesis that loss of the MotAB stator leads to increased polar localization of the MotCD stator set. Interestingly, the percentage of Δ*bifA* Δ*motAB* cells with polar localization of GFP-MotD is significantly higher than even that of WT cells. This may indicate that MotAB and MotCD do indeed compete for motor occupancy.

**Figure 1.**
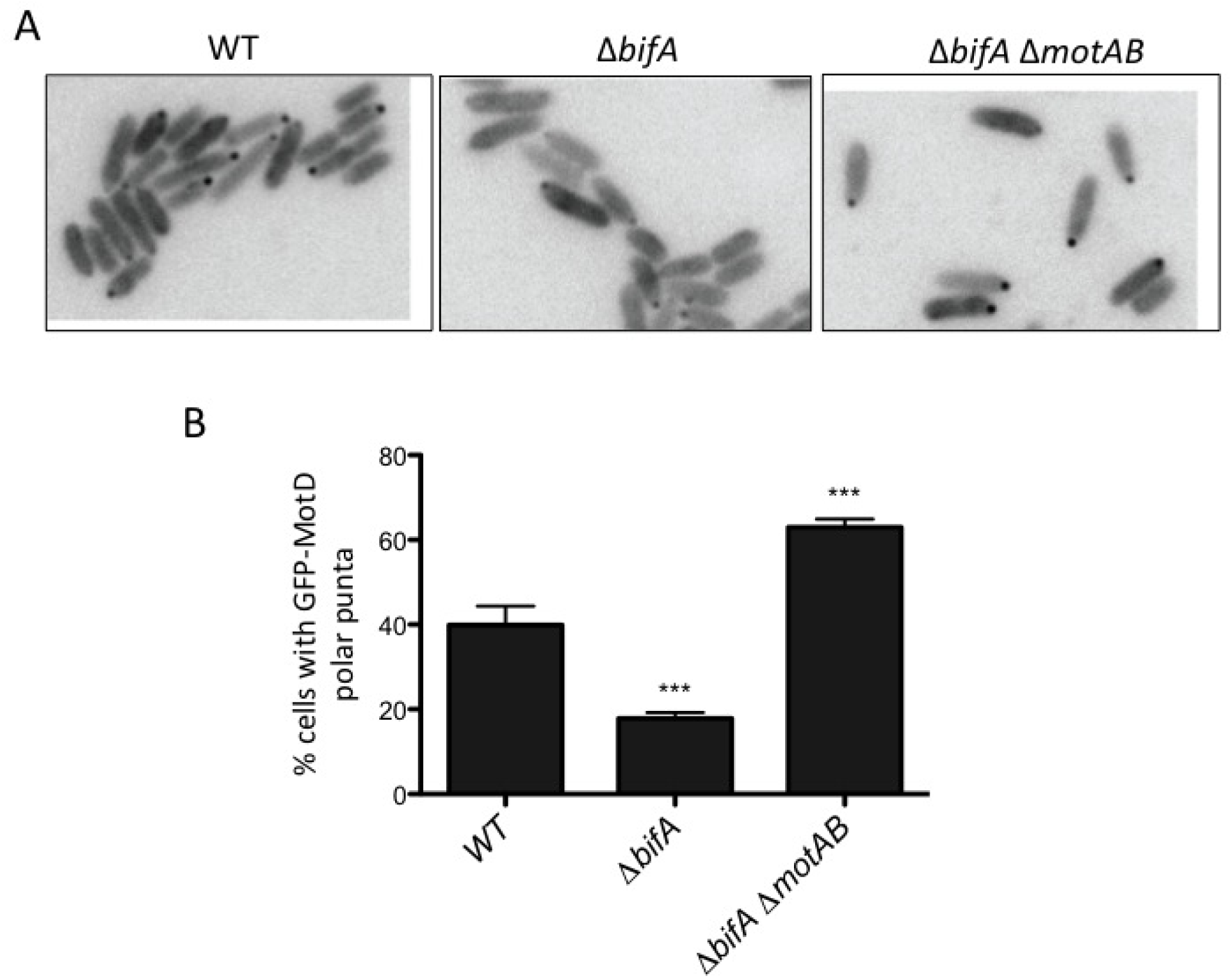
Localization of GFP-MotD is impacted by MotAB. (A) Representative fluorescence microscopy images of the indicated strains expressing GFP-MotD from the chromosome. Grayscale images were inverted using ImageJ. (B) Percentage of cells with GFP-MotD polar puncta for the indicated strains expressing GFP-MotD. These data are from two independent experiments with at least 200 total cells counted per strain per experiment. Values are reported as mean ± SEM. Significance was determined by analysis of variance and Dunnett’s posttest for comparison for differences relative to WT. ***, P < 0.001.

### Loss of stators impacts intracellular c-di-GMP levels

It is possible that increased polar localization of MotCD, rather than low c-di-GMP levels, explains the ability of the Δ*bifA* Δ*motAB* mutant to swarm (8) (Figure 2A). Thus, we hypothesized that c-di-GMP levels would remain elevated in the Δ*bifA* Δ*motAB* mutant despite its increased motility. To test this hypothesis, we used mass spectrometry to quantify levels of intracellular c-di-GMP in stator mutants. Unexpectedly, we found that c-di-GMP levels significantly decrease in the Δ*bifA* Δ*motAB* mutant relative to the Δ*bifA* mutant (Figure 2B). We also observe that the Δ*motAB* mutant shows lower cdGMP than WT, and that although in our analysis this change does not appear significant, as the level is already quite low, this lack of significance is likely the result of statistical tests having difficulty when comparing values close to zero. Together, these findings indicate that loss of MotAB is associated with decreased c-di-GMP production, particularly when levels of this dinucleotide are high.

**Figure 2.**
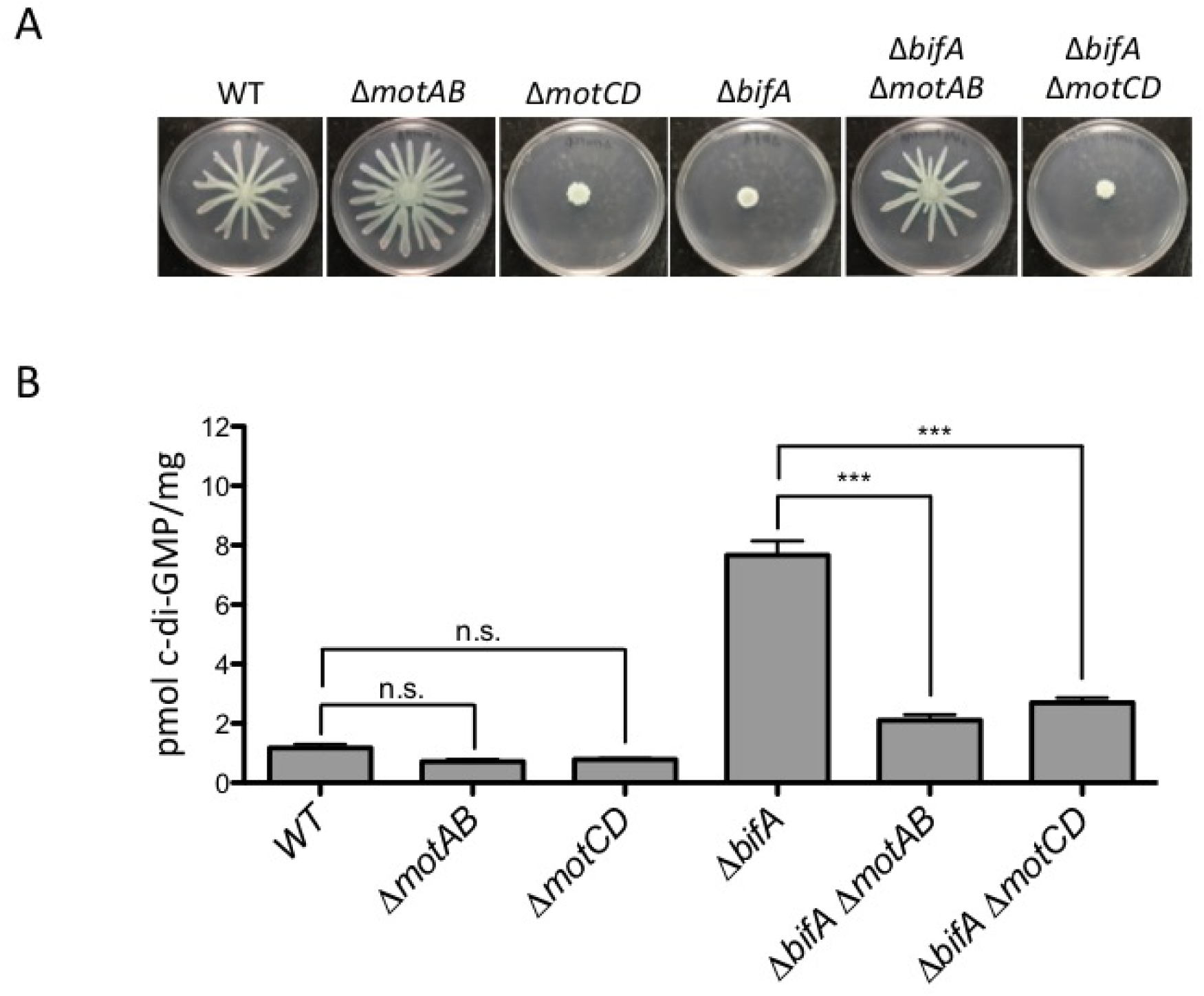
Loss of stators impacts cellular c-di-GMP levels. (A) Representative swarm plates of the strains indicated. (B) Quantification of cellular c-di-GMP levels by LC-MS for the indicated strains grown on swarm plates. Data are expressed as picomoles of c-di-GMP per mg (dry weight) of cells from which nucleotides were extracted. The data represent three independent experiments each with three biological replicates. Values are reported as mean ± SEM. Significance was determined by analysis of variance and Tukey’s posttest comparison for differences between strains indicated. n.s., not significant; ***, P < 0.001. As previously reported, Δ*bifA* is significantly different from WT (P < 0.001).

We also asked whether the “swarming-powering stator set”, MotCD affects c-di-GMP levels. We found that the Δ*bifA* Δ*motCD* mutant also has a significantly smaller pool of c-di-GMP compared to the Δ*bifA* mutant (Figure 2B). Notably, the Δ*bifA* Δ*motCD* mutant lacks the powering stator and therefore is non-motile despite having lower levels of c-di-GMP (Figure 2A,B). The Δ*motCD* mutant also has a non-significant decrease in c-di-GMP relative to WT.

Together, these results suggest that the localization and/or activities of stator proteins in the motor can impact the production of c-di-GMP. This unexpected finding links the two stator sets of *P. aeruginosa* to c-di-GMP levels of the cell.

### MotC interacts with diguanylate cyclase SadC

The effects of stator mutations on c-di-GMP levels led us to hypothesize that one or both stator sets could be interacting directly with c-di-GMP enzymatic machinery. We used a bacterial adenylate cyclase two-hybrid (B2H) assay in *E. coli* to probe for interactions between stator proteins (MotA, MotC) and two diguanylate cyclases important for swarming motility regulation, SadC and RoeA. Full-length proteins were fused to either the T18 or T25 subunit of adenylate cyclase, and then co-expressed in *E. coli* BTH101 to test for interaction. An interaction between hybrid proteins was detected as blue colored colonies on medium containing X-Gal, in which a deeper blue indicates a stronger interaction. As shown in Figure 3A, we observed an interaction between MotC and SadC, and a weaker interaction between MotA and SadC. Neither MotA nor MotC interacted with RoeA, indicating that the MotC-SadC interaction is specific. The strength of these interactions was quantified using β- galactosidase assays, and only strains co-expressing SadC and MotC fusion proteins showed significantly higher β-galactosidase than the negative control (Figure 3B).

**Figure 3.**
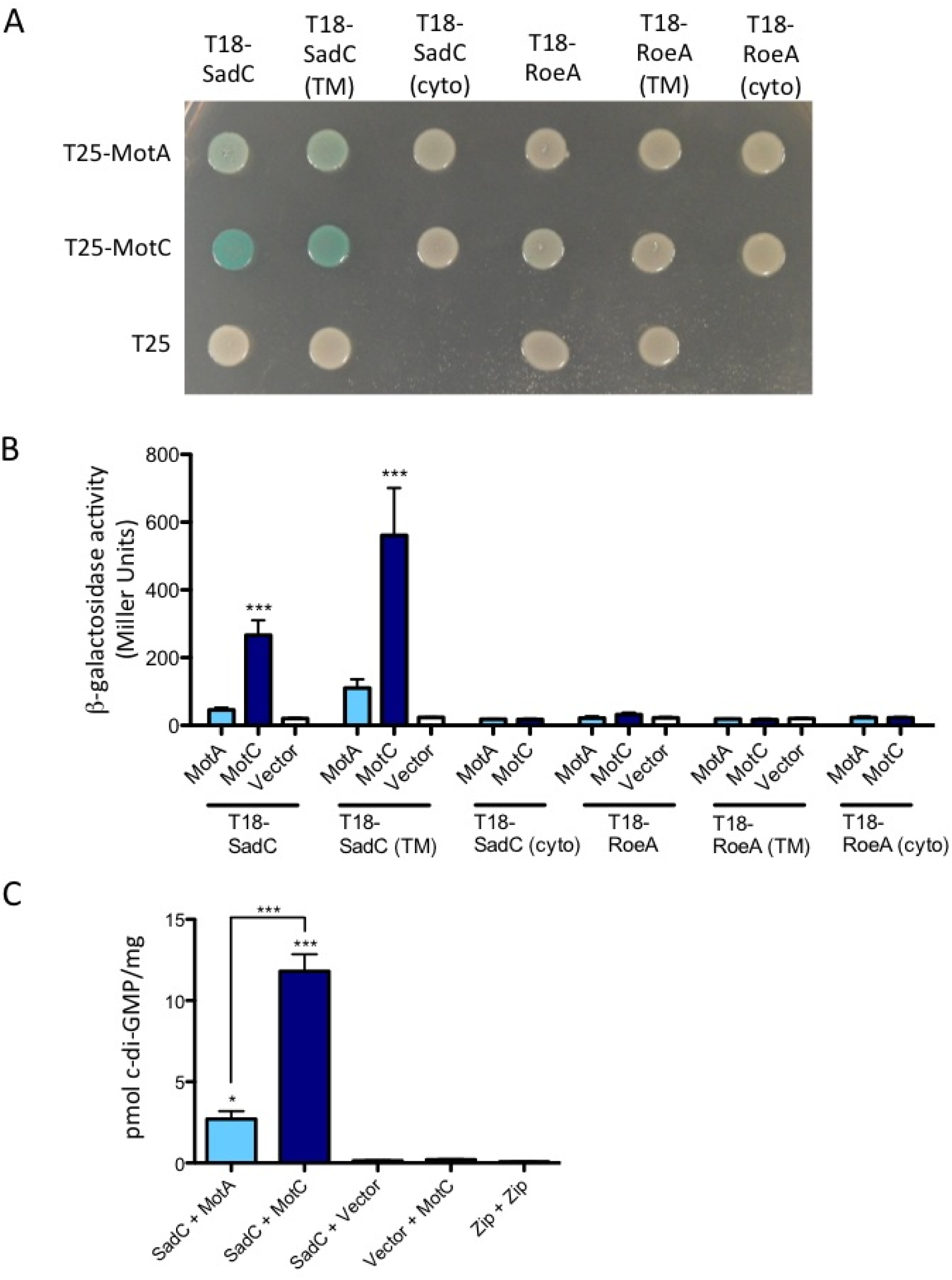
Detection of physical interaction between MotC and SadC by bacterial two-hybrid analysis. (A) The *sadC* and *roeA* genes (and truncated versions) were cloned into vector pUT18C. “TM” indicates predicted transmembrane domain and “Cyto” indicates predicted cytoplasmic region. The *motA* and *motC* genes were cloned into vector pKT25. Plasmids were co-transformed into *E. coli* BTH101 cells. The transformants were spotted (2 μl) onto LB agar containing Cb, Kan, IPTG, and X-Gal. Plates were incubated at 30°C for 40 h. Cleavage of X-Gal (blue color) indicates a positive protein-protein interaction. (B) Bacterial two-hybrid interactions were quantified by measuring β-galactosidase activity of transformants grown at 30°C overnight in LB broth supplemented with Cb and Kan. Vector indicates an empty pKT25 vector. The data represent 2 experiments each with 2-5 replicates. Values are reported as means ± SEM. Significance was determined by analysis of variance and Dunnett’s posttest comparison for difference relative to the negative control (T18-SadC + Vector). ***, P < 0.001. (C) Quantification of cellular c-di-GMP levels by LC-MS from B2H assays. The x-axis displays the two fusion proteins (listed, in order, as pUT18C + pKT25) co-transformed into BTH101 cells. After being co-transformed with 2 fusion plasmids, cells were spotted onto LB agar containing Cb, Kan, and IPTG (without X-gal). Plates were incubated at 30°C for 40 h, and then cells were scraped off plates and nucleotides were extracted. Data are expressed as picomoles of c-di-GMP per mg (dry weight) of cells from which nucleotides were extracted. The data represent 2 experiments each performed in triplicate. Values are reported as means ± SEM. Significance was determined by analysis of variance and Tukey’s posttest comparison for differences relative to the negative control (T18-SadC + Vector) and differences between the indicated samples. *, P < 0.05; ***, P < 0.001.

Both SadC and RoeA have a N-terminal transmembrane region and a C-terminal cytoplasmic domain. To identify which region of SadC may be interacting with the stators, we tested for interaction between both stators and either the transmembrane region of SadC or the cytoplasmic domain of SadC. Our results showed that MotC interacts with the transmembrane portion of SadC, but does not interact with the cytoplasmic domain of SadC. As expected, neither MotA nor MotC interact with either the transmembrane or cytoplasmic regions of RoeA (Figure 3A,B).

### The interaction between MotC and SadC stimulates SadC’s diguanylate cyclase activity

We hypothesized that association between MotC and SadC may occur when MotCD complexes are disengaged from the motor; the SadC-MotC interaction would then serve as a means to stimulate SadC activity. To test this hypothesis, we quantified c-di-GMP levels in *E. coli* BTH101 cells expressing two-hybrid constructs; we used this heterologous system because the background level of c-di-GMP produced is low and we can isolate the c-di-GMP produced by SadC from the ~30 other candidate DGCs in *P. aeruginosa*. We found that co-expression of SadC and MotC resulted in a significant increase in c-di-GMP level compared to cells co-expressing MotA and SadC (Figure 3C). These data suggest that MotC’s interaction with SadC stimulates SadC activity.

Importantly, negative controls for the B2H assay (SadC + Vector, Vector + MotC) produce relatively low levels of c-di-GMP (Figure 3C). The positive control for the B2H assay (Zip + Zip), which shows robust interaction (not shown), also produces low levels of c-di-GMP, indicating again that positive bacterial two-hybrid interactions do not generally stimulate c-di-GMP production (Figure 3C).

### Residues of the N-terminal transmembrane domain contribute to the SadC-MotC interaction and c-di-GMP stimulation

We next used the bacterial two-hybrid assay to screen for mutants of SadC (TM) with decreased interaction with MotC in order to learn more about the interaction interface between these two proteins. We used mutagenic PCR to generate mutant libraries of the T18-SadC (TM) B2H vector. We then co-transformed these libraries with a wild-type version of MotC and screened for light blue or white colonies indicating a deficiency in interaction. These mutant T18-SadC (TM) plasmids were isolated and their mutations were mapped by sequencing. A list of mutations found in this screen is presented in Supplementary Table S1.

Mutations were found throughout all of SadC’s six predicted transmembrane domains. It is likely that some of these mutations disrupt the overall structure or function of SadC along with interaction with MotC. We chose four mutations (L82P, L94P, L134R, F136Y) to introduce into the full-length SadC construct and reexamined their ability to interact with MotC. The L82P, L94P, and L134R mutants are all deficient in their ability to interact with MotC (Figure 4A,B). On the other hand, F136Y shows interaction with MotC that is comparable to wild-type SadC. It is possible that the F136Y mutation had a larger effect on interaction in the SadC (TM) construct. Alternatively, it may be a false positive from our initial screen.

**Figure 4.**
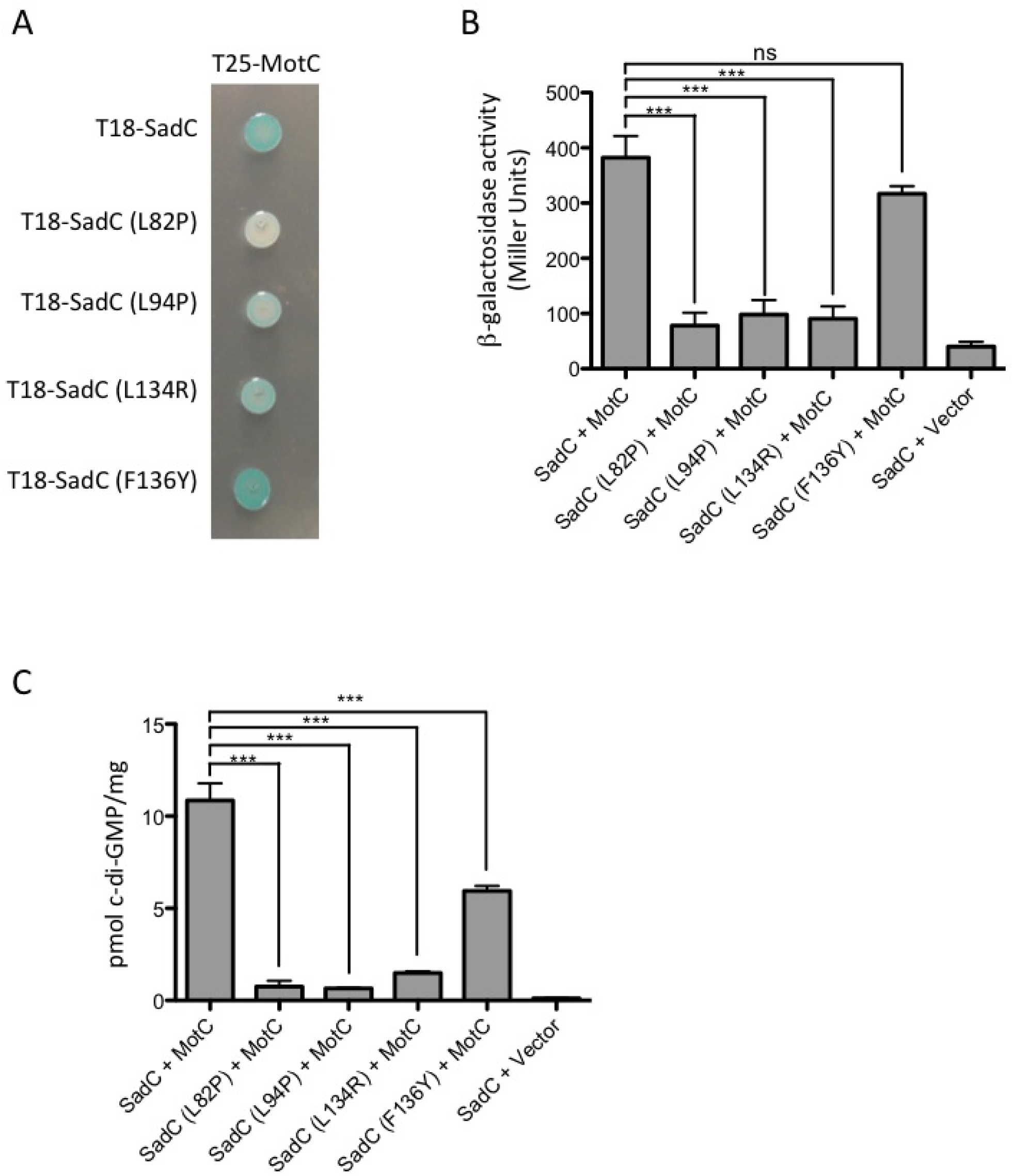
Point mutations in the transmembrane domain of SadC disrupt both its interaction with MotC and c-di-GMP production. (A) The wild-type *sadC* gene and point mutant variants were cloned into vector pUT18C. The *motC* gene was cloned into vector pKT25. Plasmids were co-transformed into *E. coli* BTH101 cells. The transformants were spotted (2 μl) onto LB agar containing Cb, Kan, IPTG, and X-Gal. Plates were incubated at 30°C for 30 h. Cleavage of X-Gal (blue color) indicates positive protein-protein interaction. (B) Bacterial two-hybrid interactions were quantified by measuring β-galactosidase activity of transformants grown at 30°C overnight in LB broth supplemented with Cb and Kan. Vector indicates an empty pKT25 vector. The data represent two experiments each performed in triplicate. Values are reported as means ± SEM. Significance was determined by analysis of variance and Dunnett’s posttest comparison for difference relative to the wild-type interaction (T18-SadC + T25-MotC). ***, P < 0.001; ns, not significant. (C) Quantification of cellular c-di-GMP levels by LC-MS from B2H assays. The x-axis displays the two fusion proteins (listed, in order, as pUT18C + pKT25) co-transformed into BTH101 cells. After being co-transformed with 2 fusion plasmids, cells were spotted onto LB agar containing Cb, Kan, and IPTG (with no X-gal). Plates were incubated at 30°C for 40 h, and then cells were scraped off plates and nucleotides were extracted. Data are expressed as picomoles of c-di-GMP per mg (dry weight) of cells from which nucleotides were extracted. Data represent three experiments each performed in triplicate. Values are reported as means ± SEM. Significance was determined by analysis of variance and Tukey’s posttest comparison for differences between strains indicated. ***, P < 0.001.

We next quantified c-di-GMP levels in BTH101 cells co-expressing the selected SadC point mutants (L82P, L94P, L134R, or F136Y) and MotC. We found that co-expression of MotC with each of the four SadC mutants resulted in a significant decrease in c-di-GMP production when compared to co-expression of MotC with wild-type SadC (Figure 4C), although the magnitude of decrease for the F136Y mutant was markedly less than the other three mutants, indicating that the F136 residue is likely not contributing robustly to the SadC-MotC interaction. We also observed that the relative strength of the SadC-MotC interaction determined in the B2H assay (Figure 4B) correlates with c-di-GMP production (Figure 4C). These results suggest that mutations that weaken SadC’s interaction with MotC lead to decreased c-di-GMP production by SadC.

### Mutants that disrupt the SadC-MotC interaction also disrupt SadC dimerization

We next asked whether any of these mutations in SadC impacted the ability of the protein to dimerize, given that DGCs like SadC function as dimers. We found that both L82P and L134R mutants showed reduced interaction with wild-type SadC (Figure 5A) and are unable to homodimerize (Figure 5B). L82P and L134R mutants were also unable to interact with MotC (Figure 4A). In contrast, both L94P and F136Y mutants are able to interact with wild-type SadC (Figure 5A), and show only a modest change in homodimerization (Figure 5B). It is interesting that these mutants have a similar ability to dimerize, given that they differ in their ability to interact with MotC – F136Y interacts with MotC, while L94P does not (Figure 4A).

**Figure 5.**
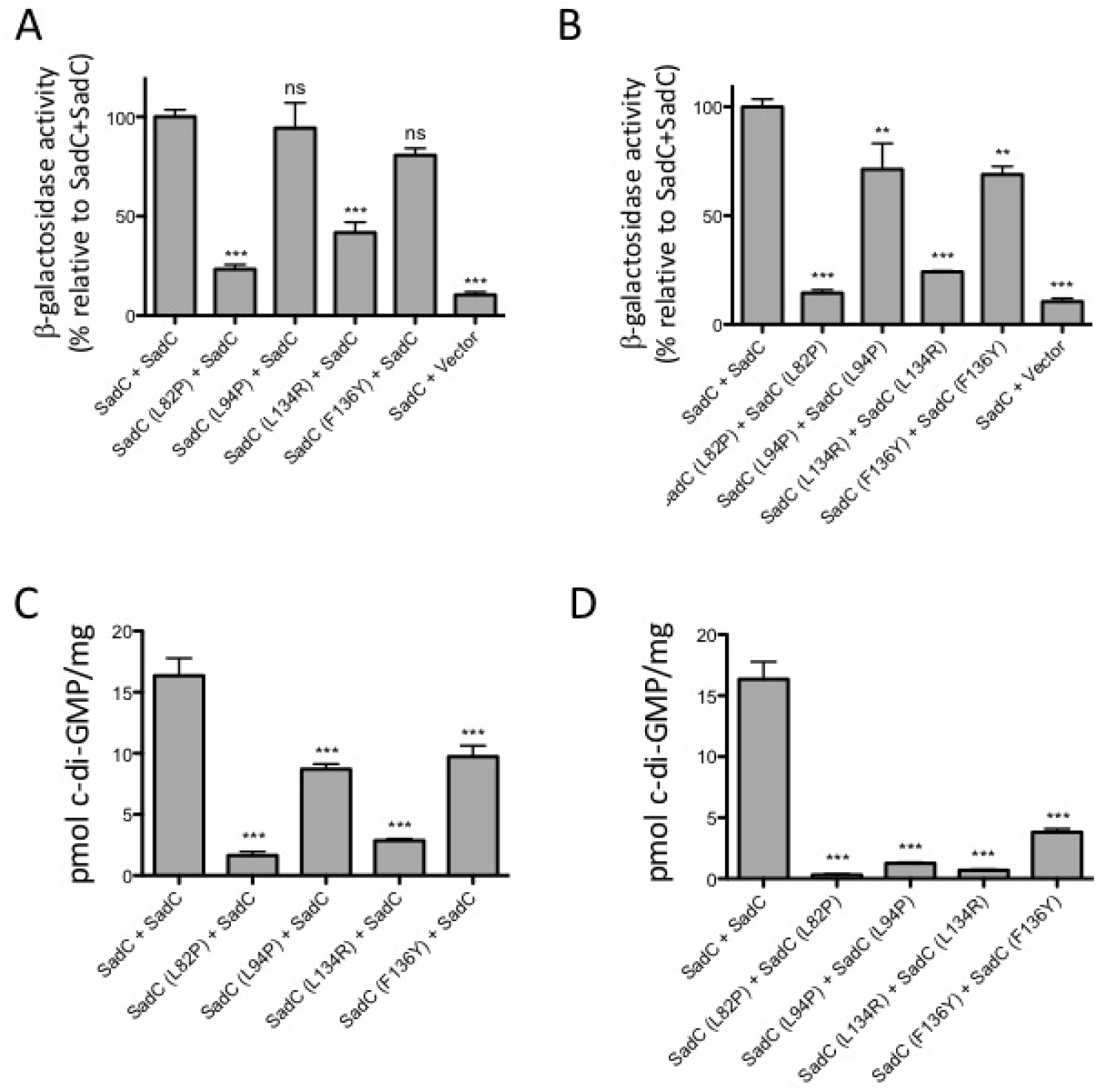
TM mutations in SadC impact dimerization and c-di-GMP production. **(A,B)** The wild-type *sadC* gene was cloned into vectors pUT18C and pKT25, and mutant *sadC* plasmids were generated using *in vitro* site-directed mutagenesis. Plasmids were co-transformed into *E. coli* BTH101 cells. The transformants were spotted (2 μl) onto LB agar containing Cb, Kan, and IPTG. Plates were incubated at 30°C for 30 h. Bacterial two-hybrid interactions were quantified by measuring β-galactosidase activity in transformants grown at 30°C overnight in LB broth supplemented with Cb and Kan. Vector indicates an empty pKT25 vector. Calculated Miller Units are represented as a percentage of the SadC + SadC interaction (set at 100%). The data from panel C and D are from 2 experiments each with 3 replicates. Values are reported as means ± SEM. Significance was determined by analysis of variance and Dunnett’s posttest comparison for difference relative to the SadC + SadC interaction. **, P < 0.01; ***, P < 0.001; ns, not significant. (C,D) Quantification of cellular c-di-GMP levels by LC-MS from B2H assays. The x-axis displays the two fusion proteins (listed, in order, as pUT18C + pKT25) co-transformed into BTH101 cells. After being co-transformed with 2 fusion plasmids, cells were spotted onto LB agar containing Cb, Kan, and IPTG (with no X-gal). Plates were incubated at 30°C for 40 h, and then cells were scraped off plates and nucleotides were extracted. Data are expressed as picomoles of c-di-GMP per mg (dry weight) of cells from which nucleotides were extracted. Data in panels E and F are from three experiments each performed in triplicate. Values are reported as means ± SEM. Significance was determined by analysis of variance and Dunnett’s posttest comparison for differences relative to the wild-type interaction (T18-SadC + T25-SadC).***, P < 0.001.

We next asked whether these SadC mutants could produce c-di-GMP as part of a SadC dimer. We found that co-expression of wild-type SadC with itself produces high levels of c-di-GMP (Figure 5C). Co-expression of wild-type SadC with any mutant SadC yielded significantly less c-di-GMP (Figure 5C). However, co-expression of SadC and SadC (L94P) or SadC (F136Y) still produced modest levels of c-di-GMP (Figure 5C). This result suggests that the L94P and F136Y mutations are having less of a negative impact on SadC activity than the L82P and L134R mutations. Overall, the L94P mutation appears to have a more specific effect on disrupting SadC’s ability to interact with MotC and to stimulate c-di-GMP production, while having only a moderate effect on SadC dimerization.

L82P and L134R homodimers produced little c-di-GMP (Figure 5D), which was expected because the physical interaction between these dimers was weak. Interestingly, although L94P and F136Y were both able to homodimerize similarly to wild-type SadC (Figure 5B,D), these mutant homodimers produced very little c-di-GMP (Figure 5D). These data indicate that physical interaction demonstrated in the B2H assay is not sufficient for c-di-GMP production; SadC may need to interact with a partner such as MotC for full activity.

### Mutations that disrupt SadC-MotC interaction alter swarming motility and reduce c-di-GMP production in P. aeruginosa

We selected two of the mutants described above (L94P and L82P) to analyze in *P. aeruginosa* to determine whether defects in the SadC-MotC interaction in the *E. coli* B2H system resulted in altered function in the native context of *P. aeruginosa*. These mutant variants of SadC were introduced onto the chromosome in the context of a C-terminal, FLAG-tagged variant of SadC. Our initial analysis revealed that the SadC-L82P mutant variant was unstable, and thus we did not work further with this mutant.

We first demonstrated that the SadC-FLAG construct conferred the same swarming phenotype as the wild-type SadC protein (Figure 6A), indicating that the C-terminal FLAG epitope did not disrupt SadC function. Next, we analyzed the impact of the SadC-L94P mutation for its ability to support swarming motility; this mutation was moderately less stable than the WT protein (the ratio of FLAG-tagged SadC/SadC-L94P = 0.79 +/-0.20, n = 4, student’s t-test, p = 0.03) (Figure 6B), but resulted in a hyper-swarming phenotype very similar to the Δ*sadC* mutant (Figure 6A). Thus, the SadC-L94P mutant protein, which does not interact with MotC, behaves as if the cell has lost all SadC function. These data are consistent with the model that SadC-MotC interaction is critical for normal swarm motility regulation.

**Figure 6.**
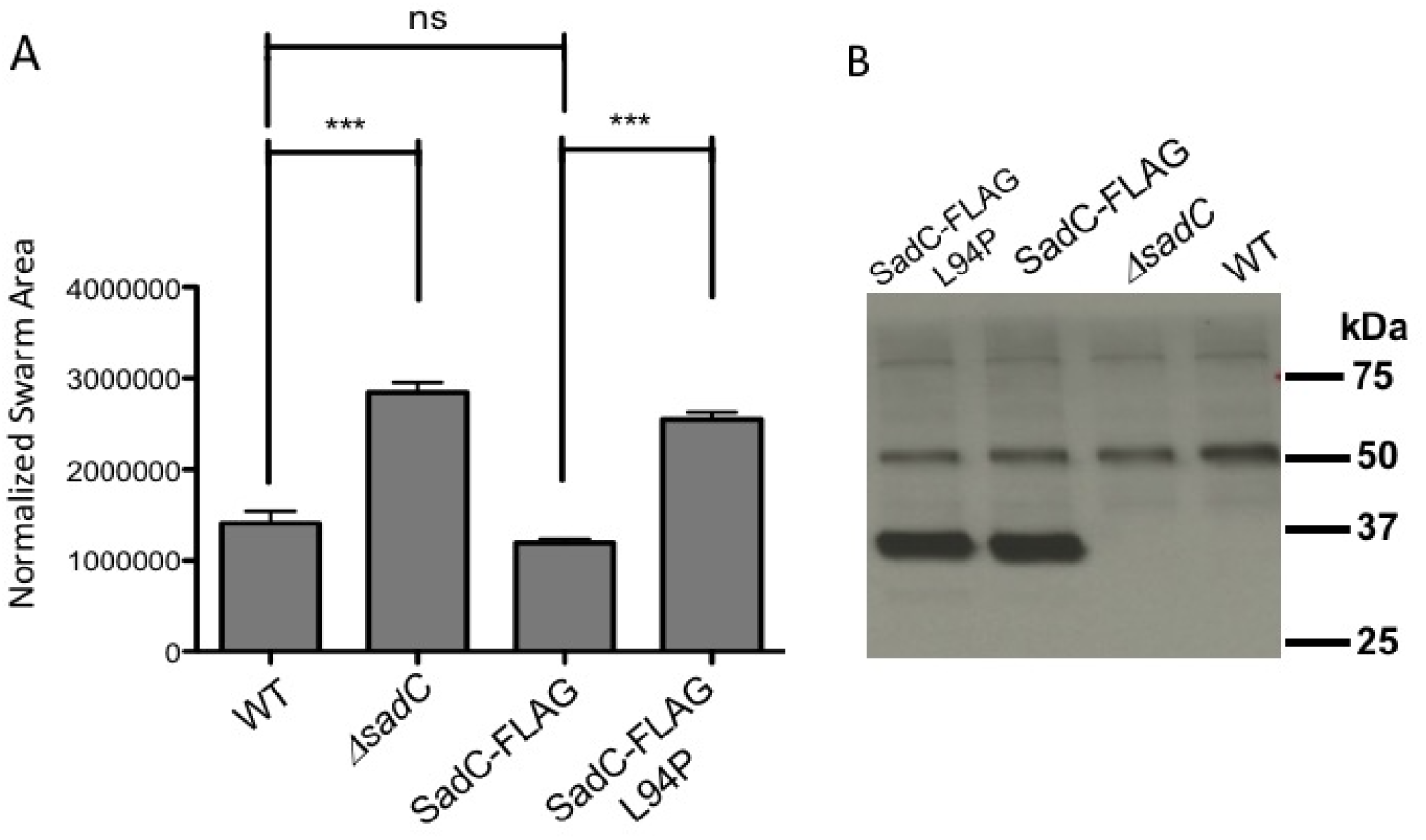
Analysis of SadC point mutations in *P. aeruginosa*. **A**. Swarming motility as assessed for the indicated strains. Significance was determined by analysis of variance and Tukey’s posttest comparison for difference relative to the WT or SadC-FLAG-expression strain swarm zone, as indicated. ***, P < 0.0001; ns, not significant. **B**. Western blot analysis of the WT and mutant FLAG-tagged SadC. The gray arrow indicates SadC. The position of the molecular weight size standards is shown on the right. The bands at ~50 and ~80 kDa are non-specific, cross-reacting bands and serve as additional loading controls. Each lane contains 250 μg of crude cell-free extract of the *P. aeruginosa* carrying the indicated allele of SadC or SadC-FLAG.

### Role of MotA in control of c-di-GMP production

We next sought to better understand the role of the MotAB stator set in the control of surface motility. We showed above that introducing a Δ*motAB* mutation into the Δ*bifA* mutant background reduces c-di-GMP levels similar to the introduction of a Δ*motCD* mutation into the same background (Figure 1). This result was surprising in that MotAB is not known to interact with SadC (Figure 3) or the c-di-GMP receptor FlgZ (9). Based on our previous studies (9) and the work above suggesting the possibility that MotAB and MotCD compete for motor occupancy, we considered the possibility that loss of MotAB would result in more MotCD in the motor, and therefore less MotCD to interact with SadC to stimulate biofilm formation.

The model proposed above suggests two strong predictions. First, if MotAB and MotCD are indeed competing for motor occupancy, and given that MotCD function is required for swarming, we would predict overexpression of MotAB to displace MotCD and thus inhibit swarming (Figure 7A, third panel). This is indeed what we observed (Figure 7B, bottom). Furthermore, the data above suggest that it is the stimulation of SadC activity by MotC interaction that drives increased c-di-GMP levels that contribute to reduced swarming. Again, our data are consistent with this model in that MotAB overexpression does not repress swarming motility in the Δ*sadC* mutant.

**Figure 7.**
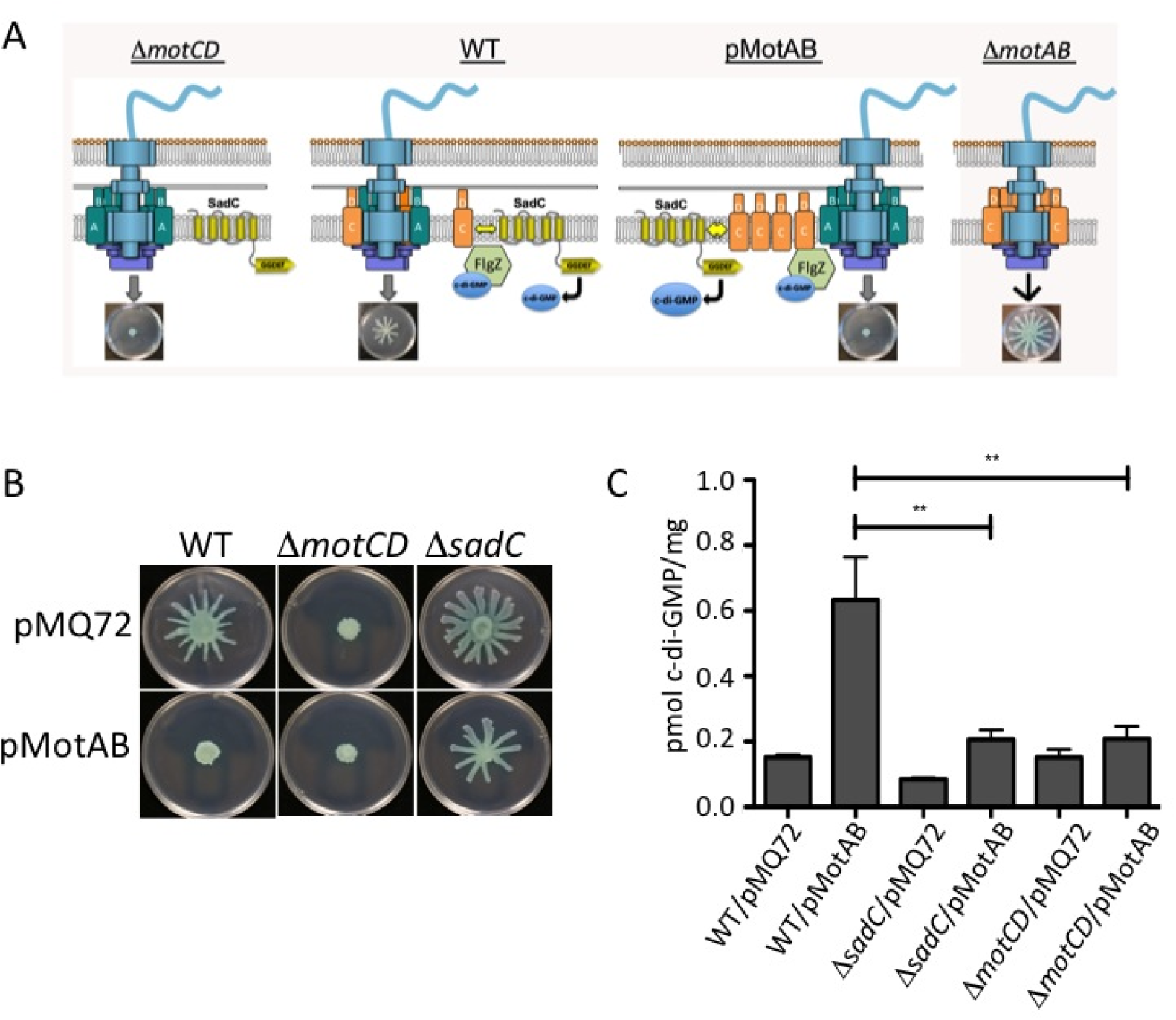
MotAB-MotCD dynamics impact c-di-GMP levels. **A.** Shown is a model of the flagellar motor with various configurations of the stators, including a Δ*motCD* mutant (left), the wild type (center, left), a strain expressing MotAB from a multi-copy plasmid (center, right), and the Δ*motAB* mutant (left). Also shown is the c-di-GMP effector FlgZ, which can interact with MotC when FlgZ is in the c-di-GMP bound state. **B**. Swarming motility as assessed for the indicated strains. **C**. c-di-GMP quantification assays. Data are expressed as picomoles of c-di-GMP per mg (dry weight) of cells from which nucleotides were extracted. Values are reported as means ± SEM. Significance was determined by analysis of variance and Dunnett’s posttest comparison to the WT carrying the vector plasmid (pMQ72). **, P < 0.01.

Second, if MotAB were indeed displacing MotCD from the stator, we would predict there would be more MotCD available to interact with SadC, and thus an increase in c-di-GMP level. We would further predict that the overexpression of MotAB would require both MotCD and SadC to increase levels of c-di-GMP. As shown in Figure 7C, overexpression of MotAB from a plasmid in the wild-type *P. aeruginosa* results in a ~5-fold increase in c-di-GMP compared to the vector control. Furthermore, this stimulation in c-di-GMP levels is dependent on the presence of SadC and the MotCD stator. Thus, it appears that MotAB can indirectly impact c-di-GMP levels via impacting the availability of MotCD to interact with SadC and thus modulate c-di-GMP production.

## Discussion

Regulation of flagellar motility via c-di-GMP is well documented and occurs at multiple levels: transcriptionally, post-transcriptionally, and post-translationally (2,12). Here we present a new kind of regulation – we show that flagellar motor components can influence c-di-GMP production. It is established that stators are dynamic, and engage and disengage from the motor in response to environmental factors such as load on the flagellum and ion availability (13). Our results indicate that the status of stators in the motor may be communicated directly to the rest of the cell through c-di-GMP signaling.

We show that deleting *motAB* in the Δ*bifA* mutant results in increased incorporation of MotCD at the motor as well as decreased c-di-GMP production. Furthermore, we found that MotC interacts with SadC in a bacterial two-hybrid assay, and that this interaction stimulates c-di-GMP production. These results may explain our finding that the Δ*bifA* Δ*motCD* mutant produces low levels of c-di-GMP; that is, the loss of MotC means this stator component cannot stimulate SadC activity.

We found that just the transmembrane domain of SadC was able to interact with MotC in the B2H assay. Recent work in *P. aeruginosa* PAO1 demonstrated that while overexpression of full-length SadC inhibits swimming motility, a truncated version of SadC lacking its transmembrane domains cannot suppress motility (even though it can still synthesize c-di-GMP) (14). Our results support these findings that membrane association is important for SadC activity, and perhaps for the ability of SadC to interact with the stator components.

Using a B2H screen, we identified point mutations in the transmembrane domain of SadC that disrupt interaction with MotC. We retested a small subset of these mutations and found that three mutations (L82P, L94P, and L134R) decrease SadC’s ability to interact with MotC and to stimulate c-di-GMP production when co-expressed with MotC in *E. coli* BTH101 cells. We further tested these SadC mutants for their ability to dimerize and the ability of those dimers to produce c-di-GMP in the context of the B2H assay. Two of these mutations (L82P and L134R) had a decreased ability to form homodimers and “heterodimers” with wild-type SadC. For these mutants, decreased dimerization led to decreased production of c-di-GMP in the B2H system.

Thus, interpreting these mutants is somewhat complicated by their apparent impact on SadC dimerization. Nevertheless, the L94P mutant could still form a dimer producing ~50% of the c-di-GMP detected for the WT SadC, and when introduced into the chromosome of *P. aeruginosa*, despite the fact that the SadC-L94P mutant protein was as stable as the WT, the L94P mutation behaved like a *sadC* null allele. Thus, we argue that this mutation’s inability to effectively interact with MotC is, at least in part, responsible for its loss-of-function phenotype.

In previous work, we proposed that MotCD disengages from the motor through its interaction with FlgZ:c-di-GMP as intracellular c-di-GMP levels increase (9). Based on results presented here, we propose that as MotCD becomes disengaged from the motor it is able to bind SadC (see Figure 7A for a model). This MotC-SadC interaction stimulates SadC activity, and thus would serve as a positive feedback mechanism to enhance c-di-GMP production when the motility-promoting stator is not in the motor. This mechanism would allow cells to directly reduce flagellar motility through stator rearrangement while further promoting sessile growth by stimulating other c-di-GMP-dependent biofilm functions (e.g., exopolysaccharide production). Furthermore, we observed that loss of the MotAB stator also resulted in reduced c-di-GMP levels. We propose that this is an indirect effect resulting from increased MotCD associating with the motor (and thus not with SadC) in the absence of MotAB, resulting in lower levels of SadC activity. Together, these data indicate a dynamic interaction among the MotAB and MotCD stators (and perhaps a competition for motor occupancy), with the SadC DGC, which allows the status of motor occupancy by the stators to be communicated to the c-di-GMP network of the cell.

## Materials and Methods

### Strains and media

Bacterial strains used in this study are listed in Table S2 in the supplementary material. *P. aeruginosa* PA14 and *E. coli* S17-1 λpir and BTH101 cells were routinely cultured in lysogeny broth (LB) or on 1.5% agar LB plates. When antibiotic selection was required, gentamicin (Gm) was used at 30 μg/ml for *P. aeruginosa*. For *E. coli*, Gm was used at 10 μg/ml, carbenicillin (Cb) at 50 μg/ml, kanamycin (Kan) at 50 μg/ml, and nalidixic acid at 20 μg/ml.

*Saccharomyces cerevisiae* strain InSc1 (Invitrogen) was used for plasmid construction using *in vivo* homologous recombination (15). InSc1 was grown in yeast extract-peptone-dextrose (1% Bacto yeast extract, 2% Bacto peptone, and 2% dextrose). Synthetic defined medium lacking uracil was used when selecting for plasmid-harboring yeast.

### Construction of mutant strains and plasmids

Plasmids used in this study are listed in Supplementary Table S3, and primers used in plasmid and mutant construction are listed in Supplementary Table S4. In-frame deletion mutants were generated using allelic exchange as previously described (15). Genes were cloned into bacterial two-hybrid vectors pKT25 and pUT18C by ligation. Point mutations made in the *sadC-*containing plasmids were constructed using a modified protocol for *in vitro* site-directed mutagenesis. Forward and reverse oligonucleotide primers were designed to contain mismatches for generating the point mutation of interest. These primers were used separately to amplify the parental plasmid using Phusion polymerase (NEB). After four amplification cycles, products of these reactions were combined and amplified for an additional 18 cycles. The parental plasmid was digested using *Dpn*I prior to transforming the PCR products into *E. coli*. Plasmids containing the desired mutation were identified by DNA sequencing.

### Swarming motility assays

Swarming assays were performed as previously described (16). Swarm medium contained M8 medium supplemented with glucose (0.2%), casamino acids (0.5%), MgSO_4_ (1mM), and 0.53% agar. Arabinose was used at 0.2% where indicated to induce expression of genes under control of the P_BAD_ promoter. Swarm plates were incubated for 16-18 h at 37°C.

### c-di-GMP measurements

*P. aeruginosa* cells were collected from swarm plates after incubation at 37°C for 18 h. For bacterial two-hybrid assay, BTH101 cells were collected from LB agar plates after 40 h at 30°C. Nucleotide extraction was performed as previously described (17, 18). C-di-GMP quantification was performed using liquid-chromatography mass spectrometry (LC-MS/MS) at the Mass Spectrometry Facility at Michigan State University.

### Fluorescence microscopy and data processing

*P. aeruginosa* for fluorescence images were picked from the edges of swarm plate colonies, as described in (16) and (8), resuspended in motility buffer (M8 medium with 0.2% glucose, 0.5% casamino acids, 1mM MgSO_4_) and 1.5 μl of the resuspension were placed on a microscope slide layered with a pad of 1.5% agarose in motility buffer. Fluorescence microscopy was performed as previously described (19) with minor modifications: a Deltavision Spectris optical sectioning microscope (Applied Precision) equipped with a UPlanApo 100 ° 1.35 oil objective (Olympus), resulting in a pixel size of 64.8 nm. A CoolSNAP_HQ2 camera (Photometrics) was used to take differential interference contrast (DIC) and fluorescence photomicrographs. For fluorophore visualization, a GFP filter set was used. DIC frames were taken with 0.01 s and fluorescence frames with 1.0 s exposure time. For each image, an initial z-stack containing 20 frames with a spacing of 150 nm was acquired. DIC frames corresponding to the center of the bacterium were selected, and 20 subsequent GFP fluorescence images at a z position corresponding to the center of the bacterium were taken. To select for stable fluorescent foci while reducing the unspecific background fluorescence present in the first two frames, frames 3-16 were averaged and background-corrected with the ImageJ software (20). The number of foci in 493-718 bacteria per strain was determined by eye by two blinded observers.

### Bacterial two-hybrid assays

Protein-protein interactions were assessed by the bacterial adenylate cyclase two-hybrid (BATCH) system obtained from Euromedex (Souffelweyersheim, France) as previously described (21, 22). Proteins of interest were fused to the T18 or T25 fragment of *Bordetella pertussis* adenylate cyclase. T18 and T25 fusion proteins were then co-expressed in *E. coli* BTH101 to test for interaction. Interaction between the two hybrid proteins reconstitutes the catalytic domain of adenylate cyclase, leading to cAMP synthesis and transcription of the *lac* operon.

BTH101 transformants were 10-fold serially diluted and spotted (2 μl) on LB agar containing Cb, Kn, X-Gal (40 μg/ml), and IPTG (0.5 mM) and incubated at 30°C for 24-40 h. The efficiencies of these interactions were quantified using β-galactosidase activity assays, as previously described (23).

### Western blot analysis

The indicated strains (WT, Δ*sadC*, and chromosomal FLAG-tagged SadC) were grown overnight in LB at 37°C and diluted 1:100 in liquid swarm medium [M8 medium supplemented with glucose (0.2%), casamino acids (0.5%) and MgSO_4_ (1mM)], and subcultured for ~3 h to OD_600_ of ~0.6. Cells were normalized to OD_600_ = 2 and lysed with sample buffer. Samples were separated on a 12% Tris-HCl gel and blotted onto nitrocellulose membrane. The membrane was incubated in blocking buffer (5% milk powder) for 1 h at room temperature. Proteins were detected with 1:10,000 dilution of monoclonal anti-FLAG M2 antibody (Sigma).

## Supporting information

## Acknowledgements

This work was supported by NIH grant R37 AI83256-06 to G.A.O. We also thank Anna Townley from the University of Oxford for assistance with analysis of fluorescence microscopy experiments and Lijun Chen at the Michigan State University Mass Spectrometry Facility for quantitative analysis of c-di-GMP.

