## Supplementary material for "Flagellar stators stimulate c-di-GMP production by *Pseudomonas aeruginosa*"

**Supplementary Table S1. SadC (TM) mutants defective for interaction with MotC isolated in a bacterial two-hybrid screen.**

| **Plasmid isolate** | **Point mutation*** | **Predicted location of mutation in full length SadC** |
| --- | --- | --- |
| 1 | **L82P** | TM3 |
| 2 | **L94P** | TM3 |
| 3 | **L134R** | TM5 |
| 4 | L29P  L148M | TM1  Between TM5 and TM6 |
| 5 | L32Q  V59E | TM1  TM2 |
| 6 | W36R  L152P | TM1  TM5 |
| 7 | L66R  F136S | TM2  TM5 |
| 8 | W67R  F139L | TM2  TM5 |
| 9 | P85A  S140P | TM3  TM5 |
| 10 | L116P  **F136Y** | TM4  TM5 |
| 11 | G22D  L57Q  I91T | TM1  TM2  TM3 |
| 12 | L37Q  A133V | TM1  TM5 |
| 13 | L66P | TM2 |
| 14 | Q86H | TM3 |
| 15 | L94Q  F166Y | TM3  TM6 |
| 16 | Y112C | TM4 |
| 17 | L117P | TM4 |
| 18 | L30P  W171G | TM1  TM6 |
| 19 | R20Q  L63P | Before TM1  TM2 |
| 20 | F121L  S140P | TM4  TM5 |
| 21 | L98P | TM3 |
| 22 | L13P  A133D | Before TM1  TM5 |
| 23 | Y44F  L134P | TM1  TM5 |
| 24 | D79G | Between TM2 and TM3 |
| 25 | Y44N  T49S  A90V  I142F | TM1  Between TM1 and TM2  TM3  TM5 |
| 26 | L104Q  R130H  I142T | TM4  TM5  TM5 |
| 27 | Y77F  L114Q  L134Q  S140P | Between TM2 and TM3  TM4  TM5  TM5 |

*Bolded alleles were retested for interaction with MotC and other phenotypes as described in the text.

**Supplementary Table S2. Strains used in this study.**

| Strain Name | Genotype/Description | Source |
| --- | --- | --- |
| ***E. coli* strains** |  |  |
| S17-1 λpir | *thi pro hsdR*- *hsdM*+ *∆recA* RP4-2::TcMu-Km::Tn7 | (1) |
| BTH101 | *F^-^, cya^-99^, araD139, galE15, galK16, rpsL1 (Str^r^), hsdR2, mcrA1, mcrB1* | Euromedex |
| S17-1 λpir | pmq30, SadC-3xFLAG KI construct, Gm^R^ | This study |
| S17-1 λpir | pmq30, SadC-3xFLAG L94P KI construct, Gm^R^ | This study |
| ***P. aeruginosa* strains** |  |  |
| SMC 232 | PA14 wild type (WT) | (Rahme et al., 1995) |
| SMC 6365 | WT (*gfp-motD*) | (2) |
| SMC 6366 | ∆*bifA* (*gfp-motD*) | (3) |
| SMC 7562 | ∆*bifA* ∆*motAB* (*gfp-motD*) | This study |
| SMC 3351 | ∆*bifA* | (4) |
| SMC 5770 | ∆*bifA* ∆*motAB* | (3) |
| SMC 5769 | ∆*motAB* | (3) |
| SMC 5684 | ∆*motCD* | (3) |
| SMC 7563 | ∆*bifA* ∆*motCD* | This study |
| SMC 8238 | SadC-3xFLAG | This study |
| SMC 8239 | SadC-3xFLAG L94P | This study |
| SMC 8240 | WT pMotAB | This study |
| SMC 7659 | ∆*motCD* pmq72 empty vector | This study |
| SMC 8241 | ∆*motCD* pMotAB | This study |
| SMC 4045 | *∆sadC* pmq72 empty vector | (5) |
| SMC 8242 | *∆sadC* pMotAB | This study |

**Supplementary Table S3. Plasmids used in this study.**

| Plasmid Name | Description | Source |
| --- | --- | --- |
| pMQ30 | Shuttle vector for yeast cloning and Gram-negative allelic replacement, Gm^r^ | (6) |
| pmq30 | SadC-3xFLAG cloned into pmq30, Gm^R^ | This study |
| pmq30 | SadC-3xFLAG L94P cloned into pmq30, Gm^R^ | This study |
| pMQ72 | Shuttle vector for yeast cloning and arabinose-inducible gene expression, Gm^r^ | (6) |
| pKT25 | BACTH vector allowing fusion to the C-terminus of the *cyaA* T25 fragment, Kan^r^ | Euromedex |
| pKNT25 | BACTH vector allowing fusion to the N-terminus of the *cyaA* T25 fragment, Kan^r^ | Euromedex |
| pUT18 | BACTH vector allowing fusion to the N-terminus of the *cyaA* T18 fragment, Amp^r^ | Euromedex |
| pUT18C | BACTH vector allowing fusion to the C-terminus of the *cyaA* T18 fragment, Amp^r^ | Euromedex |
| pKT25-zip | Leucine zipper of GCN4 fused to T25 in pKT25, Kan^r^ | Euromedex |
| pUT18C-zip | Leucine zipper of GCN4 fused to T18 in pUT18C, Amp^r^ | Euromedex |
| pUT18C-*sadC* | Full length *sadC* cloned into pUT18C, Amp^r^ | This study |
| pUT18C-*roeA* | Full length *roeA* cloned into pUT18C, Amp^r^ | This study |
| pKT25-*motA* | Full length *motA* with a C-terminal 6xHis tag cloned into pKT25, Kan^r^ | (7) |
| pKT25-*motC* | Full length *motC* cloned into pKT25, Kan | (7) |
| pUT18C-*sadC*(TM) | Transmembrane domain of *sadC* (amino acids 1-187) cloned into pUT18C, Amp^r^ | This study |
| pUT18C-*sadC*(cyto) | Cytoplasmic domain of *sadC* (amino acids 188-375) cloned into pUT18C, Amp^r^ | This study |
| pUT18C-*roeA*(TM) | Transmembrane domain of *roeA* (amino acids 1-197) cloned into pUT18C, Amp^r^ | This study |
| pUT18C-*roeA*(cyto) | Cytoplasmic domain of *roeA* (amino acids 1-187) cloned into pUT18C, Amp^r^ | This study |
| pUT18C-*sadC*(L82P) | *sadC*(L82P) cloned into pUT18C, Amp^r^ | This study |
| pUT18C-sadC(L94P) | *sadC*(L94P) cloned into pUT18C, Amp^r^ | This study |
| pUT18C-sadC(L134R) | *sadC*(L134R) cloned into pUT18C, Amp^r^ | This study |
| pKT25-sadC(L82P) | *sadC*(L82P) cloned into pKT25, Kan^r^ | This study |
| pKT25-sadC(L94P) | *sadC*(L94P) cloned into pKT25, Kan^r^ | This study |
| pKT25-sadC(L134R) | *sadC*(L134R) cloned into pKT25, Kan^r^ | This study |

**Supplementary Table S4. Oligonucleotide primers used in this study**.

| Primer Name | Primer Sequence* |
| --- | --- |
| B2H-sadC-F  B2H-sadC-R  SadC 1-187 BTH R  SadC 188-375 BTH F  SadC 188-375 BTH R  RoeA 1-197 BTH R  RoeA 198-398 BTH F  RoeA 198-398 BTH R  SadC L82P For  SadC L82P Rev | NNNNNNTCTAGAGATGCGCACAGACAAGCCTC  NNNNNNGGATCCTCGGCACTGGTGACCTCCCA  GGCGGGATCCTCGCGCATGCGTTGCCGCATC  GGCGTCTAGAGCAGCGCCGCTATGCCTTG  GGCGGGATCCTCGGCACTGGTGACCTCCCAG  GCGGGGATCCTCCCCGTGCCAGAGGATCAG  GGCGTCTAGAGCACGTGCGCAACCTGCGC  GGCGGGATCCTCCCGCAGGCTTTCCGCGAG  GTTACGCCGATCCCAGCCCGACCGAGCCGCAGGTGC  GCACCTGCGGCTCGGTCGGGCTGGGATCGGCGTAAC |
| SadC L94P For  SadC L94P Rev  SadC L134R For  SadC L134R Rev  SadC F136Y For  SadC F136Y Rev  B2H-MotA For  B2H-MotA Rev  (used a His tagged template for this amplification)  B2H-MotC For  B2H-MotC Rev  SadC native RBS pMQ72 F  SadC-His pMQ72 R  Linker + 3x FLAG G-Block sequence  SadC-3x FLAG KI F_1 upstream  SadC-3x FLAG KI R_2  SadC-3x FLAG KI F_3  SadC-3x FLAG KI F_4 downstream | GGTGGCGATCGCCTGGCCGACCTATTTCCTCTATCACGTC  GACGTGATAGAGGAAATAGGTCGGCCAGGCGATCGCCACC  CGCCCGCTGTGCGGCGCGGGCGTTCATCGC  GCGATGAACGCCCGCGCCGCACAGCGGGCG  CGCTGTGCGGCGCTGGCGTACATCGCTTTTTCCG  CCGGAAAAAGCGATGTACGCCAGCGCCGCACAGC  GGCGTCTAGAGATGTCAAAAATCATCGGCATCATCG  GGCG GGATCC TC **GTGGTGATGGTGGTGGTG**  GGCGTCTAGA G ATGGATGTGCTCAGCCTGGTC  GGCGGGATCC TC GTCCATGAAGCCTTGCAGC  caactctctactgtttctccatacccgtttttttgggTCTTCAGGCGGGTAATTCGAATG  taatctgtatcaggctgaaaatcttctctcatccgcctca**GTGGTGATGGTGGTGGTG**GGCACTGGTGACCTCCCAGG  GGCGGCAGCGGCGGCGGCAGCGGCGGCGACTACAAAGACCATGACGGTGATTATAAAGATCATGATATCGACTACAAAGATGACGACGATAAATAG  tgtaaaacgacggccagtgccaagcttgcatgcctgCCGATGCCCAGCTGGTTGACC  CGCTGTTTCAGAGGAGGCTTGTCTGTGCGCATGGGCTCCGTCCCGTAATGGCACCTGG  CGACTACAAAGATGACGACGATAAATAGGTGCCTGACATACGGGTCGGCGAGCGACGTC  ccatgattacgaattcgagctcggtacccggggatccTCAGGCGTGGGGCAGGAACAG |

* In primer sequences, uppercase boldface letters indicate a 6xHis tag, lowercase letters indicate sequence complementary to pMQ72, and underlined letters indicate point mutations.

**Literature Cited**.
